# A Deep Learning Approach for Learning Intrinsic Protein-RNA Binding Preferences

**DOI:** 10.1101/328633

**Authors:** Ilan Ben-Bassat, Benny Chor, Yaron Orenstein

## Abstract

**Motivation:** The complexes formed by binding of proteins to RNAs play key roles in many biological processes, such as splicing, gene expression regulation, translation, and viral replication. Understanding protein-RNA binding may thus provide important insights to the functionality and dynamics of many cellular processes. This has sparked substantial interest in exploring protein-RNA binding experimentally, and predicting it computationally. The key computational challenge is to efficiently and accurately infer RNA-binding models that will enable prediction of novel protein-RNA interactions to additional transcripts of interest.

**Results:** We developed DLPRB, a new deep neural network (DNN) approach for learning protein-RNA binding preferences and predicting novel interactions. We present two different network architectures: a convolutional neural network (CNN), and a recurrent neural network (RNN). The novelty of our network hinges upon two key aspects: (i) the joint analysis of both RNA sequence and structure, which is represented as a probability vector of different RNA structural contexts; (ii) novel features in the architecture of the networks, such as the application of RNNs to RNA-binding prediction, and the combination of hundreds of variable-length filters in the CNN. Our results in inferring accurate RNA-binding models from high-throughput *in vitro* data exhibit substantial improvements, compared to all previous approaches for protein-RNA binding prediction (both DNN and non-DNN based). A highly significant improvement is achieved for *in vitro* binding prediction, and a more modest, yet statistically significant,improvement for *in vivo* binding prediction. When incorporating experimentally-measured RNA structure compared to predicted one, the improvement on *in vivo* data increases. By visualizing the binding specificities, we can gain novel biological insights underlying the mechanism of protein RNA-binding.

**Availability:** The source code is publicly available at <u>https://github.com/ilanbb/dlprb</u>.

**Supplementary information:** Supplementary data are available at *Bioinformatics* online.

## 1 Introduction

The application of neural networks for machine learning (ML) purposes dates back to Rosenblatt’s perceptrons (Rosenblatt, 1958) and to Minsky and Papert’s celebrated book on the topic (Minsky and Papert, 1969). Classical learning methods often face difficulties in processing raw data, due to the need for manually designed features. In contrast, deep learning approaches can discover effective features directly from the data, and circumvent the labor-intensive phase of feature engineering. However, for over five decades, neural networks were not the method of choice in ML, as they were outperformed by a number of alternative approaches. The availability of powerful computer hardware with a large number of fast processors (such as GPUs), combined with abundant training data, has recently made deep neural networks (DNNs) the top performer in numerous ML applications. Notable areas of success include computer vision (Krizhevsky *et al.*, 2012; Szegedy *et al.*, 2015; Karayev *et al.*, 2013), natural language processing (Bowman *et al.*, 2015; Sutskever *et al.*, 2014), complex board games, such as GO (Silver *et al.*, 2016), and more.

In most areas mentioned above, hundreds of research projects utilizing DNNs were carried out and published. In computational biology the numbers are lower, yet are catching up (Angermueller *et al.*, 2016). Known applications include gene and splicing regulation (Leung *et al.*, 2014; Zhou and Troyanskaya, 2015; Kelley *et al.*, 2016), DNA methylation (Vidaki *et al.*, 2017), protein classification (Asgari and Mofrad, 2015), and various tasks in biological image analysis (Bar *et al.*, 2015; de Brebisson and Montana, 2015). Of specific relavance to our work, applications to protein-RNA binding prediction were also developed, *e.g.* DeepBind and iDeep (Alipanahi *et al.*, 2015; Pan and Shen, 2017).

The central role of protein-RNA binding in numerous biological contexts (König *et al.*, 2012) makes it an important area of study to both experimentalists and machine learning researchers. On the experimental side, high-throughput measurement techniques were developed, both for *in vivo* experiments, and for *in vitro* ones. The CLIP method and its derivatives measure protein-RNA binding *in vivo*, on a transcriptome-wide scale (Van Nostrand *et al.*, 2016; Darnell, 2010; Konig *et al.*, 2011; Hafner *et al.*, 2010). These measurements are adversely effected by a variety of orthogonal cellular events, resulting in non-negligible noise-to-signal ratio. As a consequence, these experiments are not accurate enough to produce reliable quantitative outcomes. Instead, they produce a binary outcome: yes (existence of a binding) or no (lack thereof). In one experiment, the bindings of one protein to each of its occupied transcripts *in vivo* is determined at a resolution of around 100 nucleotides. The complexity of the *in vivo* environment in the context of protein binding, as well as technological artifacts and experimental noise, make the learning of intrinsic protein-RNA binding preferences from such data a difficult challenge (Kishore *et al.*, 2011; Orenstein *et al.*, 2016b).

A different set of experimental techniques works *in vitro* (Cook *et al.*, 2017; Ray *et al.*, 2017; Lambert *et al.*, 2014). In one RNAcompete experiment, the bindings of one protein to around 240,000 short synthetic RNAs (30 to 40 nucleotides long) are measured. Lacking interfering cellular processes, these experiments exhibit low noise-to-signal ratio, and are accurate enough to produce good measurements of the bindings specificities, or strengths. The most comprehensive *in vitro* dataset, measured using the RNAcompete technology (Ray *et al.*, 2013), contains 244 such experiments (each one on a single protein).

The computational challenge that arises from these experimental data is to infer protein-specific RNA-binding models that will enable prediction of the binding between the given protein and a new RNA transcripts. Several methods have been developed to tackle this challenge. All the computational methods receive as input the RNA sequence. Some also receive the secondary structure of the RNA. We remark that the secondary structure is typically predicted by computational means, based on the sequence itself. For short RNA sequences, such computational prediction is known to be quite accurate (Doshi *et al.*, 2004). Datasets that include both RNA-binding and RNA structure measurements on the same cells are currently available for only two proteins Spitale *et al.* (2015).

The first computational method, MEMERIS, used expectation-maximization algorithm to look for sequence motifs in RNA regions that are more likely to be unpaired, and thus available for binding (Hiller *et al.*, 2006). RNAcontext, developed with the RNAcompete technology, learns a simple model for sequence and structure binding preferences (Kazan *et al.*, 2010). The sequence preferences are represented as a position weight matrix, namely how each position in the binding site contributes to the binding, independently of other positions. The structure preferences are represented as a vector of the preferences to each structural context. A more recent approach, GraphProt, uses a graph representation of RNA structure to find enriched local sub-graphs to model the sequence and structure binding preferences (Maticzka *et al.*, 2014). However, GraphProt takes more than seven days to run a single RNAcompete experiment (Orenstein *et al.*, 2016a). DeepBind, a new method based on deep learning, uses a convolutional neural network (CNN) to learn and predict protein-DNA and protein-RNA binding from many datasets, including RNAcompete and CLIP. It is based on the RNA sequence alone, i.e., without considering RNA structure (Alipanahi *et al.*, 2015). The most recent development and the state of the art, RCK, extends RNAcontext by using a *k*-mer based model, on both the sequence and on the structure level (Orenstein *et al.*, 2016a). It assigns a binding score to each RNA word of length *k* under each structural context, and thus can capture position-dependence inside a binding site. iDeep tackles the problem of predicting *in vivo* binding based on several data sources representing the complexity of the cellular environment. It receives protein binding preferences as part of the input (Pan and Shen, 2017), and thus solves a different problem. Deepnet-RBP learns RNA-binding preferences based on deep learning and using both RNA secondary and tertiary structures, but was designed to learn it from in vivo data only (Zhang *et al.*, 2015). Still, no study exploited the most advanced machine learning technique to learn intrinsic protein-RNA sequence and structure binding preferences from quantitative high-throughput *in vitro* data.

In this work, we employed two DNN architectures: convolutional neural networks and recurrent neural networks (RNNs). CNNs (LeCun*et al.*, 1998) are known to have good performance in analyzing spatial information. RNNs process input data in a sequential manner. They exploit temporal dependencies in the data, mostly by using either long short-term memory (LSTM) units (Hochreiter and Schmidhuber, 1997), or gated recurrent units (GRU) (Cho *et al.*, 2014). For both the *in vitro* case (predicting binding intensity) and the *in vivo* one (classification) our results exhibit significant improvement over all previous predictors on the same set of benchmark experiments. This comparison includes DeepBind, which employed CNN as well, but with much fewer filters and without taking RNA secondary structure into account. For *in vitro* data, our RNN achieved an average Pearson correlation of 0.628 (predicted vs. actual intensities), as compared to 0.46 by the state of the art RCK, and the runner up DeepBind with 0.41. For two different *in vivo* datasets, our CNN achieved a median AUC of 0.657 and 0.809. These results are better than the top performer DeepBind, with 0.648 and 0.803, respectively, and the improvement is statistically significant. When using experimentally-measured RNA structure as opposed to predicted on, *in vivo* binding prediction improves even further.

The remainder of this paper is organized as follows: section 2 describes the datasets used and the methods employed, including a description of DLPRB, the new RNN and CNN architectures we constructed. Section 3 presents the results of running our DNNs, and of visualizing specificities of binding sites. Finally, Section 4 contains some concluding remarks and open problems.

## 2 Methods

### 2.1 RNA secondary structure prediction

RNA secondary structural context profiles were predicted using a variant of RNAplfold (Lorenz *et al.*, 2011). In this variant, probabilities for four structural contexts are calculated per position: hairpin loop, inner loop, multi loop and external region (Kazan *et al.*, 2010). The probability for a position being paired is assigned, so that the total sum is 1. These probabilities, represented as five vectors, whose length is the same as the sequence length, were provided with the sequences as input to RCK, RNAcontext and our DNNs (DeepBind does not have such optional input).

### 2.2 *In vitro* binding prediction evaluation

To evaluate the performance of the algorithms for *in vitro* binding prediction, we used the RNAcompete dataset (Ray *et al.*, 2013). The dataset includes 244 experiments, each containing the binding intensities between a single protein and more than 240,000 RNA sequences. The set of sequences was designed as a union of two sets,A and B, such that each has similar 9-mer coverage. For each experiment,we trained a model on sequences from set A and predicted the intensities on set B. Performance was determined by the Pearson correlation of predicted and measured intensities of set B. Outlier intensities were clamped as done in the DeepBind study (Alipanahi *et al.*, 2015): all intensities above the 0.5 percentile were clamped to the value of the 0.5 percentile. Three methods were compared in this evaluation: RNAcontext, RCK and DeepBind, using results taken from (Orenstein *et al.*, 2016a).

### 2.3 *In vivo* binding prediction evaluation

For *in vivo* binding prediction, we used the eCLIP experiments (Van Nostrand *et al.*, 2016), whose proteins overlap the RNAcompete dataset. There are 21 proteins in the overlap between these two datasets. These proteins were covered by 36 RNAcompete experiments and by 54 eCLIP experiments, forming a set of 94 experimental pairs covering 21 different proteins. For each eCLIP experiment, the bound peaks were used as positive sequences, and regions 300nt downstream were used as controls, resulting in an equal-sized control set. The nearby regions were selected to test how well the binding model distinguishes between different regions on the same RNA transcript that are available for binding, while only one of them is bound. Structure prediction was performed using RNAplfold, together with 150 nucleotides flanking regions (as in previous studies (Maticzka *et al.*, 2014; Orenstein *et al.*, 2016a)), and only the original sequence peaks were used for prediction. Performance was gauged by area under the ROC curve, which is appropriate for balanced positive and negative sets, as in our case. Each binding model is trained on a complete RNAcompete experiment, and tested on its paired eCLIP experiment.

Similarly, we gauged the performance of our networks in predicting *in vivo* binding using an older dataset taken from the GraphProt study (Maticzka *et al.*, 2014). This dataset includes 23 CLIP experiments, where the intersection with RNAcompete data covers 10 proteins.

### 2.4 Data representation

Our deep learning networks receive two types of data as input, instead of a single one (Figure 1A). An RNA sequence of length *ℓ* is a string of *ℓ* nucleotides over the alphabet S = *{A, G, C, U}*. We encode every nucleotide as a one-hot vector of dimension *d*1 = 4. RNA structural information is encoded in a matrix *S ∈* ℝ^*d*_*2*_ *× ℓ*^ where *d*_*2*_ denotes the number of possible structural contexts. In this paper we consider *d*_2_ = 5 possible structural contexts.

**Fig. 1:**
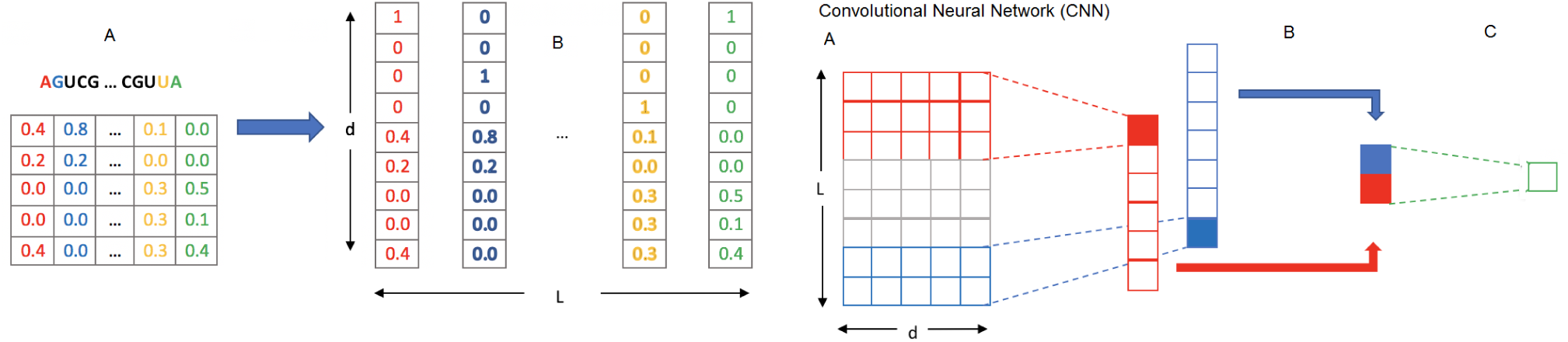
Data representation. (a) The raw input data: each sample is an RNA sequence and a matrix of structure probabilities. (b) Data transformation: each position in the sample is represented as a concatenation of two vectors: a one-hot encoding vector of the nucleotide, and a structure probabilities vector.

The information on each position in the sequence is encoded in one vector of dimension *d* = *d*_*1*_ + *d*_*2*_ (Figure 1B). Namely, every position is encoded by a concatenation of the one-hot encoding vector of the current nucleotide, and the vector of structural contexts probabilities. The method handles variable-length sequences by encoding every sequence using *L* vectors, where *L* denotes the maximum possible length of a sequence. Shorter sequences are zero-padded so that all sequences have the same length.

### 2.5 DLPRB: Convolutional and recurrent neural networks for RNA-binding prediction

The proposed CNN for predicting binding intensities receives *L*vectors for each RNA sequence, as described in section 2.4. It constructs an input matrix, *M ∈* ℝ^*L×d*^, whose rows are the vectors representing the nucleotides and structure probabilities. The input matrix, *M*, is then fed into the convolutional neural network.

Figure 2 illustrates the architecture of our CNN. The first layer of the network is a convolutional layer, which applies a series of filters on the transposed input matrices. A filter is a weight matrix *F ∈* ℝ^*m×d*^, where *d* is the dimension of the vectors representing the nucleotides and structure probabilities, and *m* is the filter length. As it is sliding, or convolving, over the input, the network computes an element-wise multiplication of the filter with all possible consecutive submatrices *W ∈* ℝ^*m×d*^ of the input data, with an addition of a bias *b*. We use a rectifier *f* (*x*) = *max*(0, *x*) as a non-linear activation function on the convolution output. The network utilizes multiple filters, with several possible values of *m*. The max-pooling layer scans the output vector of each filter and chooses the maximum value in it. A fully-connected layer computes a weighted sum of the maximum values found in the previous layer. Its output is a hidden layer of size 128. A second fully connected layer computes the final outcome of the network. We use 256 filters, 128 of length 5 and 128 of length 11. The number of filters and their lengths are different than the ones used in DeepBind.

**Fig. 2:**
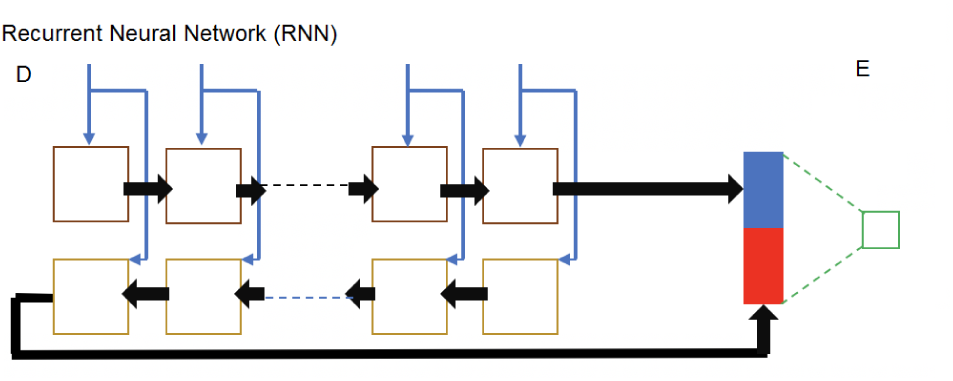
Our deep neural network architectures. (a) Convolutional layer, including a non-linear activation function. The filters are applied on a matrix whose rows are the input vectors. The output vectors of two filters are shown: one red of length three, and one blue of length two. Two specific applications of the filters are marked using dotted lines. (b) Max-pooling layer: the colored rectangles contain the max values in each vector. (c) Fully-connected layer that computes an intensity prediction. A second fully-connected layer is used in the actual implementation of the network. (d) Bidirectional recurrent neural network, composed of LSTM or GRU cells (brown rectangles). (e) Fully-connected layer that computes an intensity score. A second fully-connected layer is used in the actual implementation of the network.

Given the actual binding intensities from an RNAcompete experiment, the network is trained with a mini-batch Adam optimization algorithm (Kingma and Ba, 2014), using 128 samples in each batch, and a mean squared error (MSE) as a loss function. L2-regularization term is added to the loss function as well, by adding the sum of the squares of all the weights in the network.

Some of the hyper-parameters of the network were chosen via a grid search on a small subset of the training data. We tried several values for the number of filters (16, 64, 128 and 256), and for the filer lengths (combinations of the lengths 5, 8, 11 and 16). To reduce overfitting, the number of times the training data is processed during training (also known as the number of epochs) is tuned specifically for every dataset. This is done using a three-fold cross validation and an early-stopping procedure.

In order to assess the impact of using structure information in addition to sequence data, we predicted binding intensities with a modified CNN architecture that takes as input the one-hot encoding of the nucleotides without concatenating structure probabilities to it. This modified network was trained and tested on data that contained no structure information at all.

We also tested a bidirectional RNN, which is visually described in Figure 2. The *L* input vectors are fed into the network in a sequential manner, both forward and backward. We used GRU cells to detect possible long-term dependencies, and set the cell size to 64. The rest of the network layers, as well as the loss function and hyper-parameters tuning method are the same as in the CNN version.

### 2.6 Evaluating the weight of RNA structure in the sequence and structure binding models

To evaluate the weight of RNA structural information in our DNNs, we ran the prediction of binding while assigning uniform structural probabilities to all positions of the predicted sequences. This removes any structural information from the test data.

### 2.7 Comparing experimentally-measured and computational-predicted RNA structure

To evaluate the effect of using experimentally-measured RNA structure instead of predicted one, we used CLIP and icSHAPE data (Spitale *et al.*, 2015). Probability vectors of experimentally-measured RNA structure and CLIP-seq data were downloaded from the GEO database (accession numbers GSE60034 and GSE64168, respectively). Binding site peaks were extracted as in the original study (Spitale *et al.*, 2015) using a 40nt window size. We used the same set of peaks and control sequences as in the RCK study (Orenstein *et al.*, 2016a), summing up to 4102 sequences in each category. For computational structure prediction, we flanked binding sites and control sequences by 150nt on each end, which were only used for structure prediction by RNAplfold (Lorenz *et al.*, 2011) and later discarded for the testing.

### 2.8 Visualizing RNA-binding sequence and structure preferences

One of the main drawbacks of neural networks is their lack of interpretability. However, in some cases we can still infer how the network works, and what are the important features it extracts from the raw data. Here, we gain an understanding of how the CNN works by analyzing its filters, similarly to what was done for DeepBind (Alipanahi *et al.*, 2015). A convolution filter works like a motif detector, taking both RNA sequence and structure information into account. After training the model, we ran it to predict binding over all the test data and analyze the output of the max-pooling layer. Given a filter *F*, we extract from each test sample the subsequence to which *F* assigned the highest activation value, along with its structure information. We align all the subsequences that passed a certain threshold, and compute a modified position frequency matrix (PFM), that also captures the structure information for every position in the sequence. From this matrix we then generate the sequence and structure logos (Wagih, 2017).

## 3 Results

### 3.1 Predicting *in vitro* binding

To gauge the performance of DLPRB, our deep neural networks, compared to extant methods, we used the comprehensive dataset of RNAcompete, which includes 244 experiments (Ray *et al.*, 2013). We trained a model on one half and tested it on the other half. Performance was gauged by Pearson correlation of predicted and measured intensities. For complete details see Subsection 2.2.

Both our deep neural networks significantly outperformed all methods in *in vitro* binding prediction (Figure 3A). When comparing them to extant methods, they both outperformed the state of the art RCK, which achieved an average Pearson correlation of 0.46 as compared to 0.606 for CNN (p-value= 9.16 *·* 10^−42^, Wilcoxon rank-sum test). RNN outperformed CNN, achieving an average Pearson correlation of 0.628 (p-value=9.53*·*10^−26^).

**Fig. 3:**
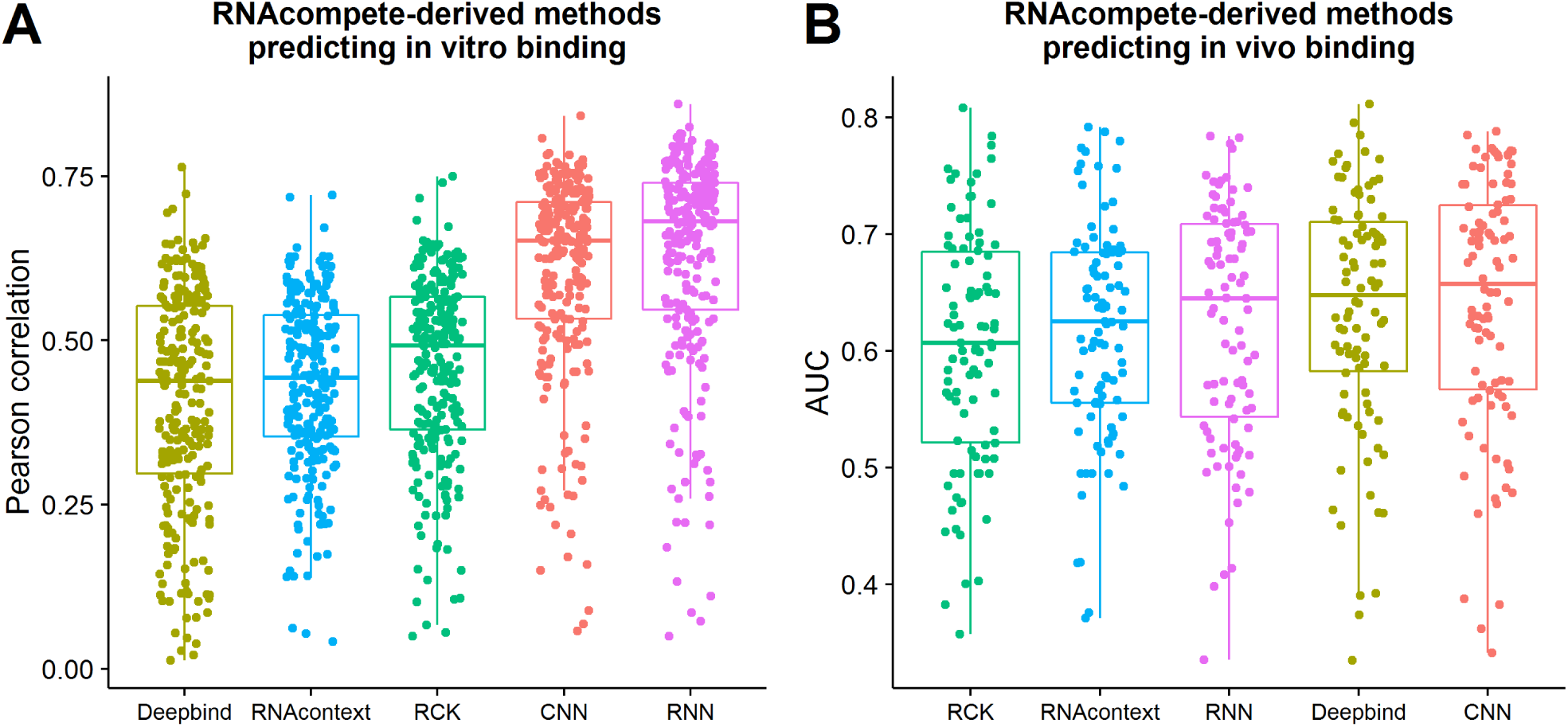
Performance of RNAcompete-derived methods in predicting binding shown as boxplots for different methods. RNAcontext, RCK, CNN and RNN utilize RNA secondary structure. (A) Performance in predicting *in vitro* binding. For each RNAcompete experiment, a model was trained on Set A sequences and tested on Set B. Results are over 244 experiments. Performance gauged by Pearson correlation of predicted and measured intensities.(B) For each RNAcompete experiment, a model was trained on the whole dataset and tested in predicting bound and unbound transcripts as measured by eCLIP experiment on the same protein. Results are over 94 experiment pairs. Performance gauged by area under the ROC curve.

When training and testing our CNN on RNA sequence alone,we saw a statistically significant difference in prediction accuracy comparing to training and testing on both RNA sequence and structure (average Pearson correlation dropped from 0.606 to 0.592, p-value=1.26 *·* 10^−24^). As was the case in(Orenstein *et al.*, 2016a), this shows that structure information can improve binding prediction. We speculate that the relatively small difference is due to the fact that RNA structure was predicted from sequence, and DNNs are capable of capturinglong-range nucleotide interactions, which are the basis of RNA secondary structure. Moreover, the library design of RNAcompete technology was designed to be unstructured, and as a consequence, contains very few structures (Ray *et al.*, 2013, 2009). Thus, it is no surprise that the sequence features alone can capture most of the structural information. Still, the statistically significant difference shows the importance of RNA structure in protein-RNA binding prediction.

As the gap between our CNN and the network used by DeepBind was not fully explained by removing RNA structural information, we ran a variant of our approach with much fewer convolution filters. Instead of using 256 filters of variable lengths, we used only 16 filters, each of length 16, imitating the configuration used in (Alipanahi *et al.*, 2015). The results dropped profoundly to an average Pearson correlation of 0.523 compared to 0.592, just by decreasing the number of filters and changing their lengths (p-value=4.69 *·* 10^−41^). We deduce that our advantage over DeepBind can be explained mainly by either the configuration of the convolution filters, or by the RNN architecture. In addition, RNA structural information also improves the accuracy of our prediction. For complete results see Supplementary Table S1.

### 3.2 Predicting *in vivo* binding

To gauge the performance of DLPRB, our neural networks,compared to extant methods on *in vivo* binding prediction, we used the eCLIP dataset (Van Nostrand *et al.*, 2016). The overlap with RNA compete dataset covers 21 proteins by 94 experimental pairs involving 36 RNAcompete and 54 eCLIP experiments (Ray *et al.*, 2013). Each binding model was trained on a complete RNAcompete experiment, and tested on its paired eCLIP experiment. We report the performance in predicting *in vivo* binding by AUC, an appropriate metric for balanced bound and unbound sets as in our case. Similarly, we test our method on an older CLIP dataset taken from the GraphProt study (Maticzka *et al.*, 2014). The overlap with RNAcompete covers 10 proteins. For complete details see Subsection 2.3.

The results of predicting *in vivo* binding show that CNN performs the best, achieving a median AUC of 0.657, compared to 0.648 and 0.645 for DeepBind and RNN, respectively (Figure 3B). In a pairwise comparison CNN is significantly better than DeepBind and RNN (p-values<0.0001, Wilcoxon rank-sum test). When tested on an older dataset, our CNN network outperformed all other methods achieving a median AUC of 0.809 compared to 0.803 and 0.782 for DeepBind and RNN, respectively (Figure 4A). This improvement is not statistically significant as there were only 10 proteins in the overlap with RNAcompete for this dataset.

**Fig. 4:**
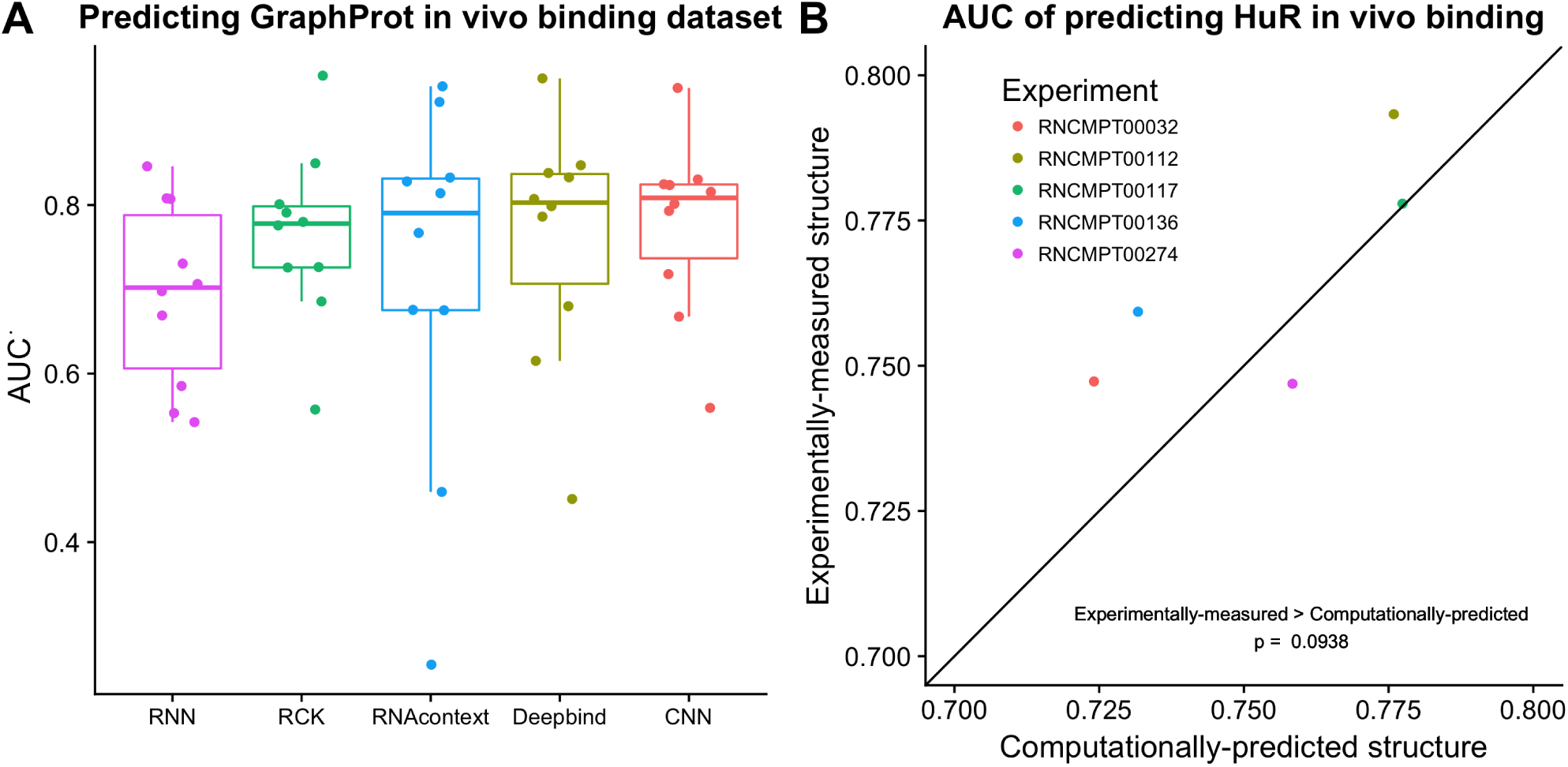
Improved *in vivo* binding prediction. A) We used GraphPror dataset of *in vivo* binding to gauge prediction accuracy. For each pair of RNAcompete and CLIP experiments on the same protein,a model was trained on the former and tested on the latter.23 pairs overlap with the GraphProt study and RNAcompete dataset, covering 10 proteins in 21 RNAcompete and 12 CLIP experiments. Performance per protein is gauged by average AUC.B) We compared experimentally-measured and computationally-predicted RNA structure. We used available CLIP and iCSHAPE datasets that had an RNAcompete experiment available for the same protein. The only dataset tested HuR protein and had five corresponding RNAcompete experiments. A model trained on RNAcompete data had better binding predicting with experimentally-measured RNA structure than computational-predicted structure.

Two reasons may hamper the accuracy of *in vitro* models in predicting *in vivo* binding. First, *in vivo* data is known to be noisy and suffer from experimental biases (Kishore *et al.*, 2011). Moreover, RNA structure prediction is less accurate *in vivo* than *in vitro* (Rouskin *et al.*, 2014), so learned structural preferences may not improve binding prediction. At this stage, more datasets with higher quality are needed in the overlap between CLIP and RNAcompete to derive more definitive conclusions. For complete results see Supplementary Table S2.

### 3.3 Experimentally-measured structure may improve *in vivo* binding prediction

Following the results in the previous subsection, we examined the reason why RNA structural information did not improve *in vivo* binding prediction. We speculated that RNA structure prediction of long RNA transcripts *in vivo* is inaccurate. To test this hypothesis, we compared computational-predicted RNA structure (Lorenz *et al.*, 2011) with experimentally-measured one (Spitale *et al.*, 2015) in the task of binding prediction. To demonstrate the effect of using experimental probabilities, we used available CLIP and icSHAPE experiments, performed on the same cells that also had an RNAcompete experiment on the same protein. Unfortunately, only the HuR protein, which had five RNAcompete experiments, was found to overlap. Since icSHAPE reports only unpaired probabilities, we trained a model based on two structural contexts: paired and unpaired. Performance was measured by AUC in predicting HuR binding sites. For complete details see Subsection 2.7.

Results show that our neural networks benefit from experimentally-measured RNA structure in predicting *in vivo* binding (Figure 4B). icSHAPE measurements are more accurate than predicted structure in four out of five experiments. We note that additional experimental measurements of RNA structure and protein-RNA binding on the same cells are needed to evaluate the benefit of experimentally-measured RNA structure for the task of *in vivo* binding prediction. For complete results see Supplementary Table S4.

### 3.4 The weight of RNA structure in the sequence and structure binding models

We gauged the weight of RNA structural preferences in binding prediction using our CNN architecture. We employed a comprehensive dataset of both *in vitro* and *in vivo* data as in previous sections. For each test set we used the same models, trained on both RNA sequence and structure data, but we now predicted binding using uniform structure probabilities, and compared them to predictions using predicted structure probabilities. As we already noted in Section 3.1, when training and testing on RNA sequence alone, the prediction accuracy is slightly, albeit significantly, lower compared to one achieved by using RNA structure on top of sequence. Despite the fact that RNAcompete sequence set was designed to be unstructured, previous studies have shown that some structure exists and that RNA structural binding preferences can still be inferred from the data (Ray *et al.*, 2013; Orenstein *et al.*, 2016a). For completed details see Subsection 2.6.

We found that the use of predicted structure probabilities in test time significantly improves prediction performance *in vitro*, but not *in vivo*. In terms of *in vitro* binding, the improvement is across the board: for every single dataset, the performance improved by using predicted structure probabilities as compared to uniform ones (Figure 5A). The improvement was striking, from an average Pearson correlation of 0.54 for sequence-only mode, to 0.608 when using predicted structure probabilities (p-value = 4.53 *·*10^−42^, Wilcoxon rank-sum test). The results of the *in vivo* data, on the other hand, did not show significant improvement (p-value = 0.96) (Figure 5B). We can see several reasons for this dichotomy. The *in vivo* dataset is smaller - only 94 pairs compared to 244, covering only 21 proteins compared to 205. The *in vivo* experiments are noisier and prone to technological artifacts (Kishore *et al.*, 2011). The *in vivo* environment contains many confounding factors, which are not part of the binding model, and thus may decrease prediction accuracy. Lastly, RNA secondary structure is less accurate for long sequences *in vivo* than for short sequences and *in vitro* (Rouskin *et al.*, 2014). For complete results see Supplementary Table S3.

**Fig. 5:**
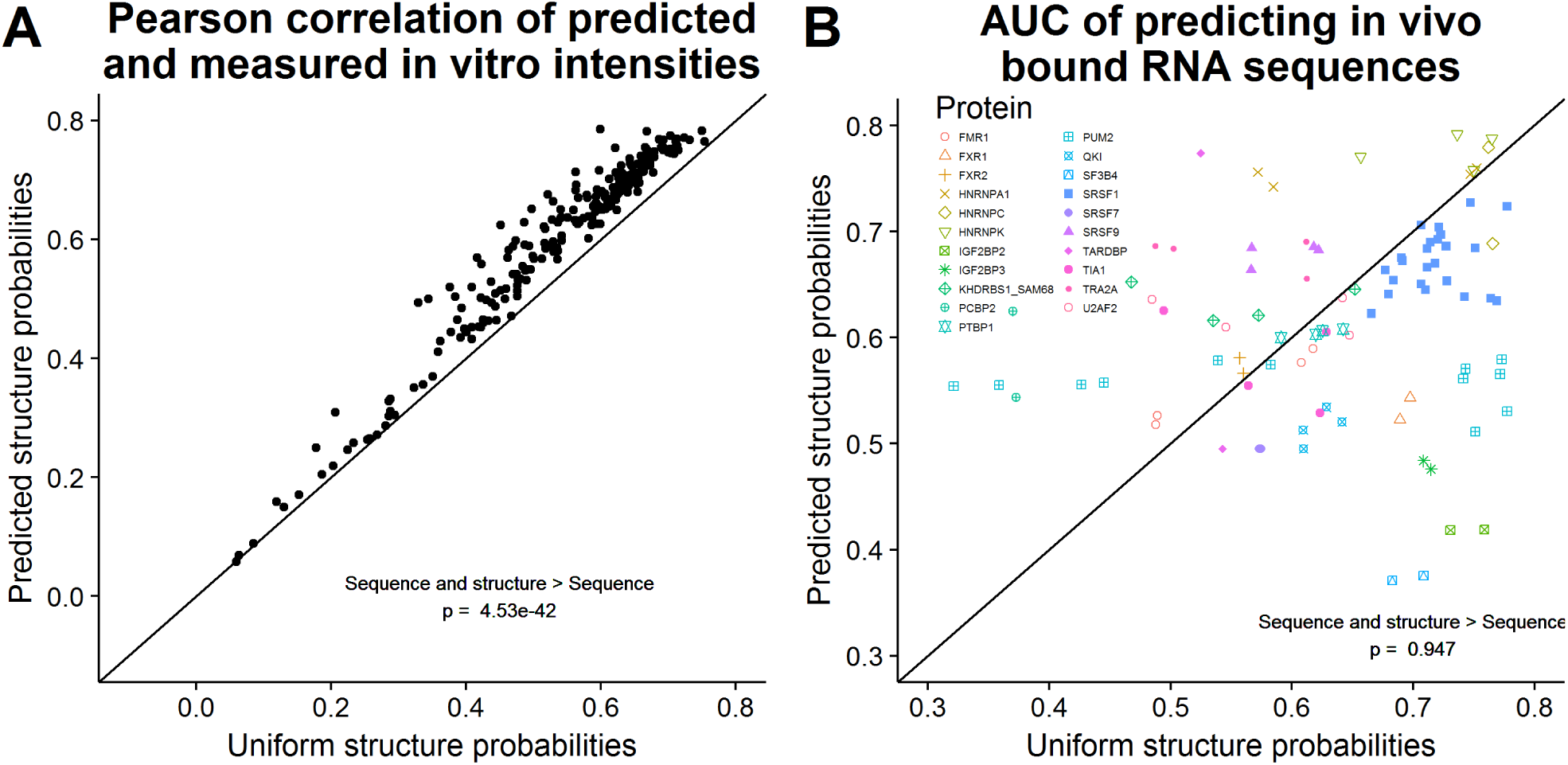
The weight of RNA structure on top of sequence in binding prediction. (A) Predicted RNA structure probabilities improve *in vitro* binding prediction compared to uniform ones. Correlation results over 244 paired experiments uncovers that RNA structure plays a significant role in protein-RNA interactions. D) Predicted RNA structure probabilities do not improve *in vivo* binding prediction compared to uniform ones. AUC results of 96 paired eCLIP and RNAcompete experiments over 21 joint proteins demonstrate that RNA structure is not accurately predicted for *in vivo* transcripts, and that protein-intrinsic binding preferences do not capture the full complexity of the cellular environment.

### 3.5 Visualizing RNA-binding specificities

Finally, we wanted to learn new biological insights on the RNA sequence and structure binding preferences of the proteins in the RNAcompete dataset. Interpreting deep convolutional neural networks is a long-standing challenge which we do not solve in this study.Instead,we developed a heuristic. We look for hits of the motif detector,i.e.,binding sites that passed a certain threshold, and use their alignment to generate a position frequency matrix for both the RNA sequence and RNA structure probabilities. We draw these as sequence logos (Wagih, 2017). For complete details see Subsection 2.8.

Figure 6 shows a comparison of sequence logos generated by different methods on different datasets for three proteins: Pum2, Vts1 and HuR. We see a high concordance in the sequence and structure preference as discovered by GraphProt (Maticzka *et al.*, 2014) for Pum2 protein trained on an independent PAR-CLIP dataset (Hafner *et al.*, 2010), and for HuR and Vts1 as discovered by RNAcontext (Kazan *et al.*, 2010) trained on an old version of RNAcompete (Ray *et al.*, 2009). This demonstrates the ability of our CNN to learn true RNA sequence and structure binding preference and to generate interpretable visualization of them.

**Fig. 6:**
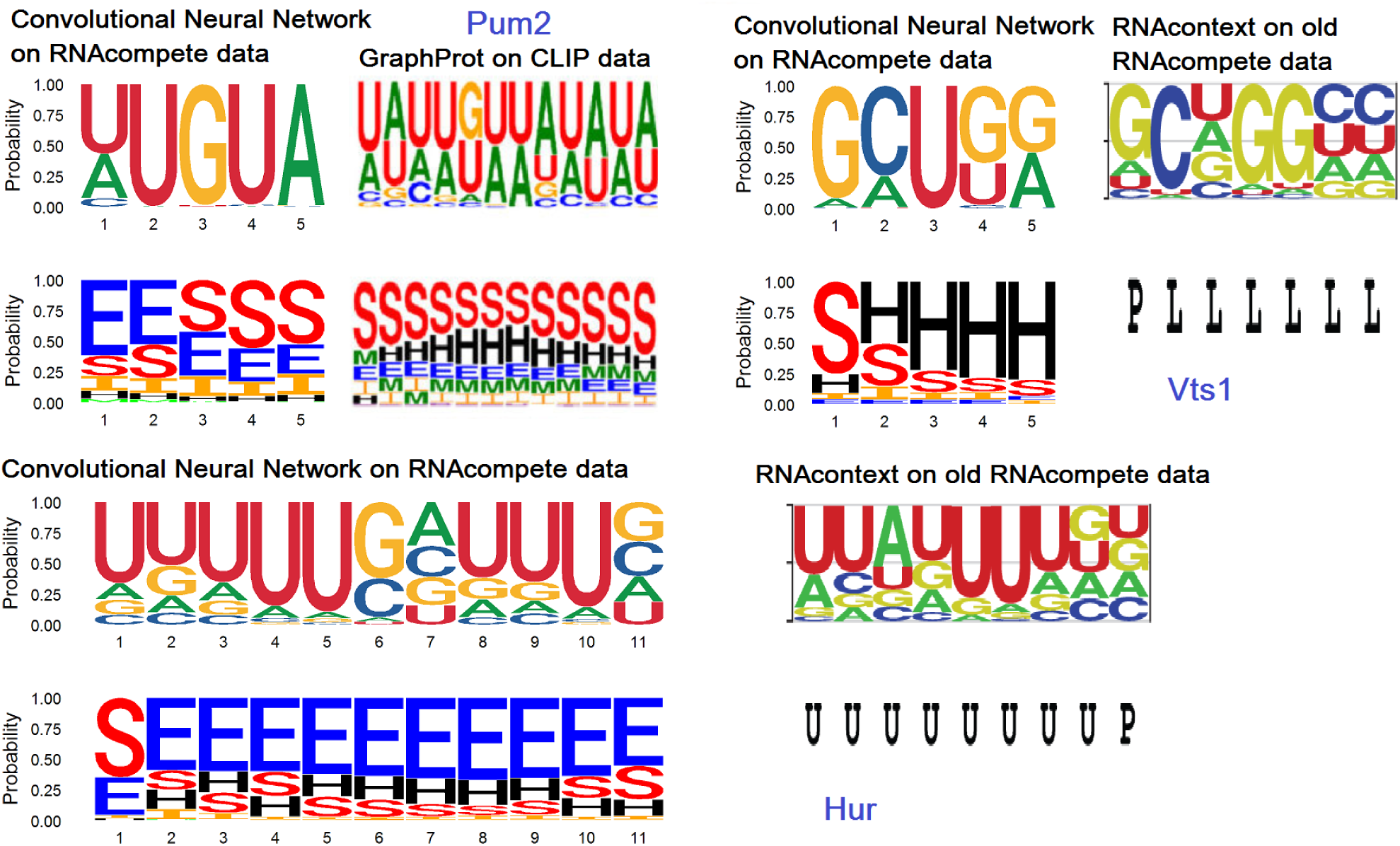
Sequence and structure binding preference visualization. We visualize the binding preferences, represented as two frequency matrices, sequence and structure, by the sequence logo format. Compared to previous methods, GraphProt (Maticzka *et al.*, 2014) and RNAcontext (Kazan *et al.*, 2010), trained on different datasets than those used in our study (Hafner *et al.*, 2010; Ray *et al.*, 2009), we see high concordance in both the sequence and structure preference. U,P,L stand for unpaired, paired and loop. S,H,I,M,E stand for stem (paired), hairpin, inner, multi and external.

## 4 Discussion

We have shown that carefully designed deep neural networks are capable of significantly improving the predictive power in protein-RNA binding experiments. By using different network architectures,and by incorporating structure information in the learning process, we outperformed the state of the art results for this task. The improvement was substantially noticeable for *in vitro* experiments. This is possibly due to the fact that *in vitro* experiments are designed to quantitatively measure protein-RNA binding for hundreds of thousands of synthetic RNA sequences. We believe these results demonstrate the usefulness of deep neural networks in the subject of protein-RNA binding, and more generally in the field of computational biology, where they are starting to be used on a large scale.

A long-standing goal in the field of protein-RNA interaction is accurate prediction of *in vivo* binding. As demonstrated in this study, current computational methods perform poorly in predicting bound and unbound RNA transcripts (average AUCs around 0.65, and some predictions are even below 0.5, which corresponds to random guessing). We believe that learning the intrinsic binding preferences of an RNA-binding protein would not suffice in this case, as the *in vivo* environment is much more complex. Not only do proteins compete over the same binding sites or co-bind together, RNA structure also differs between *in silico, in vitro* and *in vivo* environments. On top of that, RNAcompete and other *in vitro* experiments measure binding to short RNA sequences (30-40nt) (Ray *et al.*, 2009; Lambert *et al.*, 2014), which cannot fold to complex RNA structures that are found *in vivo*, where transcripts span thousands of nucleotides. This alone already inhibits *in vitro* trained models from learning binding preferences to complex structures.

There are a number of questions worth pursuing following our work: Why were RNNs better than CNNs for *in vitro* data, but worse than them for *in vivo* data? Our training and test data were based on experiments where the binding between a single protein and numerous RNAs was measured. Can we design a DNN (or another ML mechanism) to train on many proteins and RNAs, and then to predict the binding of *different* proteins and RNAs? Another future line of research is to further improve the interpretability of the suggested networks. In particular, a better understanding of how the structure information is incorporated in the learning and prediction processes, and what filters are more dominant and why, may yield interesting biological insights.

## Funding

I.B.B. was supported by fellowships from the Edmond J. Safra Center for Bioinformatics at Tel-Aviv University and the Blavatnik Research Fund.

